# Co-occurrence of Carbapenemase Genes, Disinfectant Resistance Determinants, and Class 1 Integrons in *Acinetobacter* spp. Isolated from Bloodstream Infections in Critical Care Units

**DOI:** 10.64898/2026.04.27.721256

**Authors:** Tofazzal Md Rakib, Farhan Fuad Bin Hossen, Shantu Chowdhury, Pair Ahmed Jiko, Shourav Mohajan, Sabreena Alam, Ayesha Ahmed Khan, Shohag Majumder, Md. Arafat, Nurul Absar, A M A M Zonaed Siddiki

**Affiliations:** Department of Pathology and Parasitology, Chattogram Veterinary and Animal Sciences University, Chittagong, Bangladesh 4225; Transboundary Animal Diseases Research Center, Kagoshima University, Kagoshima, Japan 8900065; Department of Clinical Biochemistry, Institute of Applied Health Sciences Hospital, Chattogram, Bangladesh; Department of Biochemistry and Biotechnology, University of Science and Technology Chittagong, Bangladesh; Department of Microbiology, Institute of Applied Health Sciences Hospital, Chattogram, Bangladesh; Department of Emergency and Critical Care Medicine, Aichi Medical University, Nagakute, Japan

**Keywords:** *Acinetobacter*, Multidrug-resistance, Carbapenem-resistance, Biocide resistance, Class integrons

## Abstract

*Acinetobacter* spp. represents critical opportunistic pathogens driving severe bloodstream infections (BSIs) in intensive care unit (ICU) and neonatal intensive care unit (NICU) settings. The convergence of carbapenem resistance and emerging biocide tolerance, often mediated by mobile genetic elements, has intensified concerns regarding co-selection and persistence in clinical environments. A total of 90 molecularly confirmed *Acinetobacter* isolates (ICU = 44; NICU = 46) from bloodstream infections were analyzed. Antimicrobial susceptibility was determined using the Kirby–Bauer disk diffusion method in accordance with CLSI M100 (2024) guidelines and extended-spectrum β-lactamase production was assessed by combined disc diffusion. Polymerase chain reaction (PCR) was employed to detect carbapenemase genes (bla_VIM_, bla_NDM_, bla_IMP_, bla_OXA-23_, bla_OXA-58_), biocide resistance determinants (*qacE*, *qac*Δ*E1*), and the class 1 integron-integrase gene (*intI1*). Multidrug-resistant (MDR) and extensively drug-resistant (XDR) phenotypes were identified in 71.1% (64/90) and 22.2% (20/90) of isolates, respectively. High resistance (>71%) was observed against meropenem and cephalosporins, whereas colistin (58.8%, 53/90) and amikacin (47.8%, 43/90) showed moderate susceptibility. The most prevalent genotypes were *qac*Δ*E1* (76.6%, 69/90) and *bla_VIM_* (56.6%, 51/90). Statistical and network analyses revealed significant correlations between biocide and carbapenemase genes, identifying *IntI1* as a primary driver of co-resistance. The findings indicate a significant co-occurrence of carbapenemase genes, biocide resistance determinants, and class 1 integrons among *Acinobacter* spp. isolates. These associations suggest that mobile genetic elements may contribute to the dissemination of resistance traits.

## 1. Introduction

Bloodstream infections (BSIs) caused by *Acinetobacter* spp. have emerged as a persistent and escalating threat in modern healthcare, with a disproportionate burden in intensive care units (ICUs) and neonatal intensive care units (NICUs), where host vulnerability, invasive procedures, and prolonged antimicrobial exposure converge [1–4]. Among these organisms, *Acinetobacter baumannii* has attained particular clinical prominence due to its extraordinary capacity for environmental persistence and rapid acquisition of resistance determinants, driving high rates of morbidity, mortality, and healthcare-associated transmission. In recognition of this threat, the World Health Organization has designated carbapenem-resistant *A. baumannii* as a critical priority pathogen, underscoring the urgent need for comprehensive surveillance and mechanistic insights into resistance evolution [5–7].

Carbapenem resistance in *Acinetobacter* spp. is predominantly mediated by the production of β-lactamases, including class D OXA-type enzymes (e.g., bla_OXA-23_, bla_OXA-24/40_, bla_OXA-58_) and class B metallo-β-lactamases (MBLs) such as bla_VIM_, bla_IMP_, and bla_NDM_ [8–12]. These determinants are frequently embedded within mobile genetic elements, notably class 1 integrons which facilitate horizontal gene transfer and accelerate the dissemination of resistance across bacterial populations. The integron-integrase gene *intI1* plays a central role in capturing and expressing gene cassettes, thereby enabling the accumulation and co-expression of multiple resistance determinants [13]. This genetic plasticity underpins the global expansion of multidrug-resistant (MDR) and extensively drug-resistant (XDR) *Acinetobacter* lineages, severely constraining therapeutic options and complicating clinical management [14–16].

Beyond antibiotic resistance, increasing attention has been directed toward the role of biocide tolerance in shaping resistance ecology. Genes such as *qacE* and its truncated derivative *qac*Δ*E1*, commonly associated with class 1 integrons, encode efflux systems that reduce susceptibility to quaternary ammonium compounds widely used in hospital disinfection [15–18]. Biocide resistance genes such as qacE and qacΔE1 are frequently reported in association with class 1 integrons and antibiotic resistance determinants. Their concurrent occurrence may facilitate co-selection under disinfectant and antimicrobial pressure [1, 5, 6].

Despite accumulating evidence on individual resistance mechanisms, there remains a critical gap in understanding the integrated landscape of phenotypic resistance, carbapenemase gene distribution, and biocide resistance determinants particularly in the context of bloodstream infections from ICUs and NICUs in resource-constrained settings. Furthermore, the extent to which mobile genetic elements such as *intI1* orchestrate the co-occurrence and interaction of these determinants has not been sufficiently elucidated.

In this context, the present study systematically investigates the antimicrobial susceptibility profiles and molecular epidemiology of *Acinetobacter* spp. isolated from bloodstream infections in ICU and NICU settings. By combining phenotypic susceptibility testing with targeted molecular screening of carbapenemase and biocide resistance genes, alongside integron-associated elements, this work aims to delineate the co-resistance architecture driving MDR and XDR phenotypes. Elucidating these relationships is essential for informing evidence-based infection control strategies, optimizing disinfection protocols, and strengthening antibiotic stewardship frameworks in high-risk clinical environments.

## 2. Materials and Methods

### 2.1. Sample Collection and Bacterial Isolation

A total of 122 blood culture specimens obtained from patients admitted to ICUs and NICUs of four tertiary hospitals in Chattogram, Bangladesh, between June 2024 and September 2025 were included. Only the first Acinetobacter-positive isolate recovered from each patient was included in the study to avoid duplicate sampling. All specimens were transported under appropriate conditions to the Department of Pathology and Parasitology, Chattogram Veterinary and Animal Sciences University (CVASU) for microbiological analysis. Written informed consent was obtained from all participants or their legal guardians prior to sample collection. The study protocol was reviewed and approved by the institutional ethics review board.

Upon receipt, blood samples were inoculated onto blood agar plates and incubated aerobically at 37 °C for 18–24 h. Colonies exhibiting characteristic morphology suggestive of *Acinetobacter* spp. were subcultured to obtain pure isolates. Preliminary identification was performed using standard biochemical assays, including triple sugar iron (TSI) agar reactions, motility testing, citrate utilization, oxidase and catalase activity, and oxidative–fermentative (OF) tests. Molecular confirmation was subsequently performed by PCR targeting the conserved *recA* gene, resulting in 90 confirmed *Acinetobacter* spp. bloodstream isolates.

### 2.3. Antimicrobial Sensitivity Testing

Antimicrobial susceptibility profiles were determined using the Kirby–Bauer disc diffusion method on Mueller–Hinton agar (Oxoid, UK), following Clinical and Laboratory Standards Institute (CLSI) guidelines [19]. Bacterial suspensions were standardized to 0.5 McFarland and incubated at 37 °C for 18–24 h. A total of 17 antibiotics representing seven antimicrobial classes were tested, including aminoglycosides (amikacin, gentamicin), cephalosporins (cefuroxime, ceftriaxone, ceftazidime, cefotaxime, cefepime), β-lactam/β-lactamase inhibitor combinations (piperacillin–tazobactam, amoxicillin–clavulanic acid), carbapenems (imipenem, meropenem), fluoroquinolones (ciprofloxacin, levofloxacin), tetracyclines (tetracycline, doxycycline), and folate pathway inhibitors (trimethoprim–sulfamethoxazole). The CLSI (2022) criteria were used to determine the breakpoints; isolates with MICs ≤2 µg/mL were considered intermediate, whereas those with MICs ≥4 µg/mL were considered resistant. *Escherichia coli* CVASU K-18 was used as a quality control strain. Classification of isolates as multidrug-resistant (MDR), extensively drug-resistant (XDR), or pan drug-resistant (PDR) was performed according to the criteria described by Asaduzzaman et al. [20].

### 2.4. Genomic DNA extraction

Genomic DNA was extracted from confirmed *Acinetobacter* isolates using a modified boiling method [21]. Briefly, 3–5 freshly cultured colonies were suspended in 100 µL of DNase-free water to achieve an approximate concentration of 1 × 10[CFU/mL. The suspension was vortexed and subjected to thermal lysis at 95 °C for 10 min, followed by centrifugation at 13,000 ×g for 5 min. The supernatant containing genomic DNA was collected and stored at −20 °C. DNA concentration and purity were assessed using a NanoDrop One spectrophotometer (Thermo Fisher Scientific, USA).

### 2.5. Molecular Detection of Resistance Genes

Detection of antimicrobial resistance genes was performed by polymerase chain reaction (PCR). Each reaction was carried out in a total volume of 25 µL containing 12.5 µL of GoTaq® Green Master Mix (Promega, USA), 1 µL each of forward and reverse primers (10 µM), 1 µL of DNA template, and nuclease-free water to volume.

PCR amplification was conducted in a FlexCycler2 thermocycler (Analytik Jena, Germany) under the following conditions: initial denaturation at 95 °C for 1 min; 35 cycles of denaturation at 95 °C for 30 s, annealing at 55–60 °C for 40 s, and extension at 72 °C for 90 s; followed by a final extension at 72 °C for 4 min. Amplified products were resolved by electrophoresis on 1.5% agarose gels stained with ethidium bromide. Band sizes were determined using a 100 bp DNA ladder (Promega, USA) and visualized under UV illumination using a UVP UVsolo Touch imaging system (Analytik Jena, Germany).

### 2.6. Phenotypic Detection of ESBL Production

Extended-spectrum β-lactamase (ESBL) production was assessed among carbapenem-resistant *Acinetobacter* isolates (n = 55) using the phenotypic combination disc diffusion test (PCDDT). Bacterial suspensions equivalent to 0.5 McFarland (approximately 1.5 × 10[CFU/mL) were inoculated onto Mueller–Hinton agar plates.

Discs containing cefotaxime (30 µg) and ceftazidime (30 µg), alone and in combination with clavulanic acid (cefotaxime/clavulanic acid 30/10 µg and ceftazidime/clavulanic acid 30/10 µg), were placed at a minimum distance of 25 mm (center-to-center). Plates were incubated at 37 °C for 18–24 h. An increase in inhibition zone diameter of ≥5 mm for combination discs compared to the corresponding single antibiotic discs was interpreted as indicative of ESBL production.

### 2.7. Statistical analysis

Statistical analyses were performed using GraphPad Prism version 10.3.2 and Python (version 3.13). Categorical variables were summarized as frequencies and percentages. Associations between categorical variables were evaluated using the Chi-square test or Fisher’s exact test, as appropriate based on expected cell counts. A p-value of ≤0.05 was considered statistically significant.

## 3. Results

### 3.1 Distribution of *Acinetobacter* spp. Isolates

Of the 122 blood samples processed, 90 isolates were confirmed as *Acinetobacter* spp. by recA-based PCR. All isolates were recovered from bloodstream infections, indicating bacteremia as the predominant clinical presentation. The distribution across critical care units was comparable, with 51.1% (46/90) originating from NICUs and 48.9% (44/90) from ICUs.

Male predominance was observed in both settings (ICU: 60.8%, NICU: 59.0%), yielding male-to-female ratios of 1.56:1 and 1.45:1, respectively (Figure 1A-B). Age-stratified analysis revealed the highest proportion of Acinetobacter-positive bloodstream infections among ICU patients aged 18–44 years (36.9%) and 45–64 years (41.3%). In contrast, NICU cases were predominantly early (≤7 days; 38.6%) and late neonates (8–28 days; 31.8%) (Table 2).

**Figure 1:**
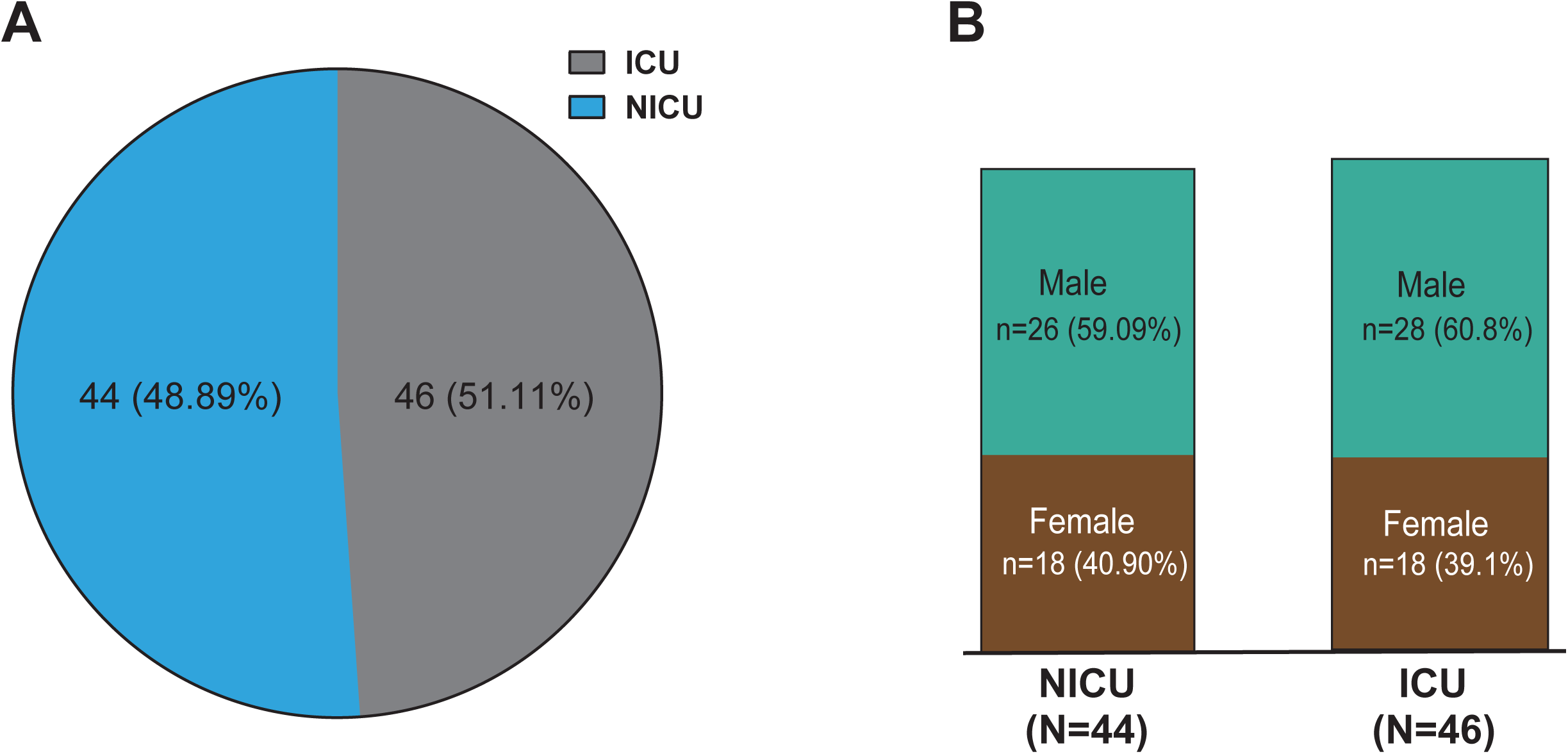
Distribution of *Acinetobacter* spp. isolates among patients in intensive care units (ICU) and neonatal intensive care units (NICU) (A), and corresponding sex distribution within each group (B).

**Table 1:**
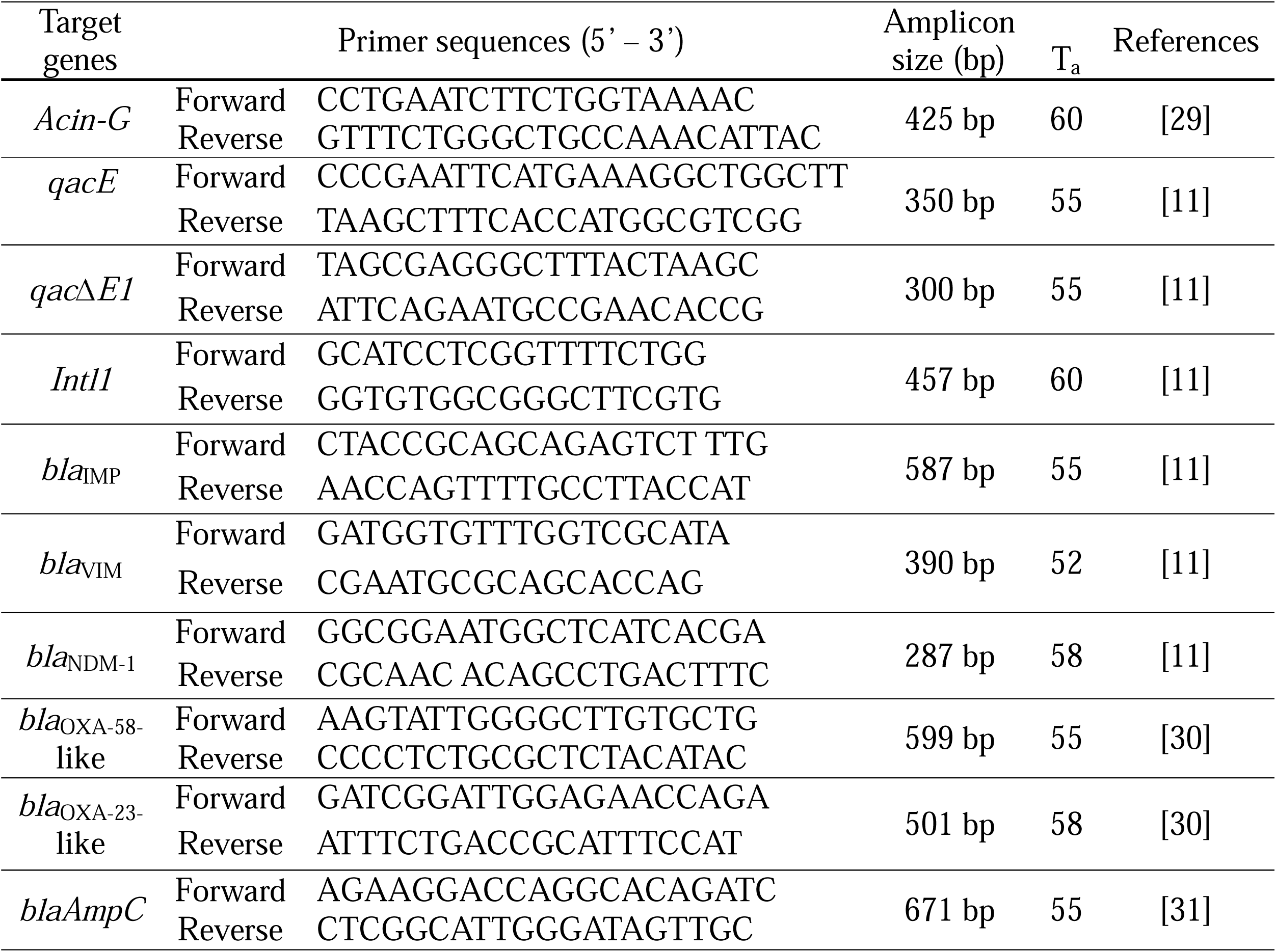
Primer sequences and amplification characteristics used for molecular detection of resistance determinants in this study.

**Table 2:** Age distribution of patients stratified by intensive care unit (ICU) and neonatal intensive care unit (NICU) cohorts.

| Age Categories |  | % |
| --- | --- | --- |
| **ICU (n=46)** |  |  |
| <17 Years (Child) | 4 | 8.70 |
| 18 – 44 Years (Young Adult) | 17 | 36.96 |
| 45 – 64 Years (Middle-Aged) | 19 | 41.30 |
| > 65 Years (Elderly/Geriatric) | 6 | 13.04 |
| **NICU (n=44)** |  |  |
| <7 days (Early Neonatal) | 17 | 38.63 |
| 8-28 days (Late Neonatal) | 14 | 31.81 |
| <1 (Infant/Adolescent) | 13 | 29.54 |

### 3.2 Antibiotic Susceptibility Profiles

Among the 90 isolates, a high burden of antimicrobial resistance was observed, with 71.1% (n=64) classified as multidrug-resistant (MDR), 22.2% (n=20) as extensively drug-resistant (XDR), and only 6.7% (n=6) as non-MDR (Figure 2A). Stratified analysis showed higher MDR prevalence in NICU isolates (75.0%) compared to ICU isolates (67.3%), whereas XDR was more frequent in ICU isolates (28.2% vs. 15.9%) (Figure 2B).

**Figure 2:**
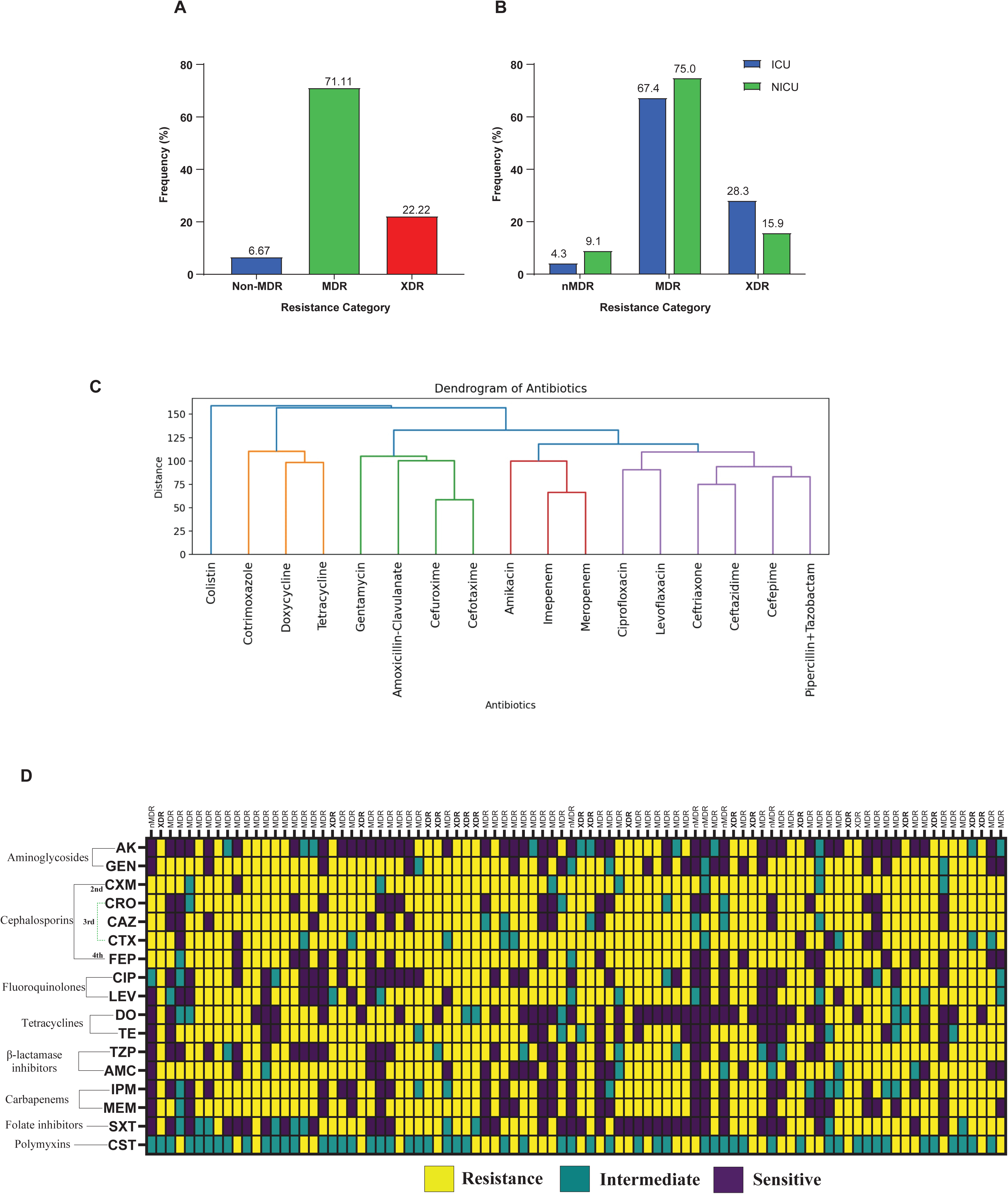
Comprehensive phenotypic characterization of antimicrobial resistance among *Acinetobacter* spp. isolates (N = 90). (A) Overall distribution of resistance phenotypes, including non-multidrug-resistant (non-MDR), multidrug-resistant (MDR), and extensively drug-resistant (XDR) isolates. (B) Comparative distribution of resistance phenotypes between intensive care unit (ICU; n = 46) and neonatal intensive care unit (NICU; n = 44) cohorts. (C) Hierarchical clustering dendrogram of isolates based on phenotypic susceptibility patterns, generated using Ward’s linkage method and distance-based metrics, demonstrating similarity and clustering of resistance profiles. (D) Heatmap illustrating antibiotic susceptibility profiles across all isolates; rows represent individual isolates stratified by resistance phenotype, while columns denote antibiotics grouped by class. Color gradients indicate susceptibility status: resistant (yellow), intermediate (green), and susceptible (purple). Here, AK, Amikacin; GEN, Gentamicin; CXM, Cefuroxime; CRO, Ceftriaxone; CAZ, Ceftazidime; CTX, Cefotaxime; FEP, Cefepime; CIP, Ciprofloxacin; LEV, Levofloxacin; DO, Doxycycline; TE, Tetracycline; TZP, Piperacillin–Tazobactam; AMC, Amoxicillin–Clavulanate; IPM, Imipenem; MEM, Meropenem; SXT, Cotrimoxazole; CST, Colistin.

Overall susceptibility profiling demonstrated extensive resistance across antibiotic classes (Table 3). Cephalosporins exhibited the highest resistance rates, including cefuroxime (91.1%), cefotaxime (83.3%), ceftazidime (78.8%), cefepime (75.5%), and ceftriaxone (74.4%). High resistance was also observed for gentamicin, levofloxacin, tetracycline, and amoxicillin–clavulanate (each 72.2%), as well as meropenem (71.1%). In contrast, relatively higher susceptibility was retained for colistin (58.8%), amikacin (47.8%), doxycycline (44.4%), and trimethoprim–sulfamethoxazole (40.0%).

**Table 3:** Overall antimicrobial susceptibility profile of *Acinetobacter* spp. isolates.

| Class | Antibiotics | S (n) | % | I (n) | % | R (n) | % |
| --- | --- | --- | --- | --- | --- | --- | --- |
| Aminoglycosides | Amikacin | 43 | 47.77 | 11 | 12.22 | 36 | 40.00 |
|  | Gentamycin | 19 | 21.11 | 6 | 6.67 | 65 | 72.22 |
| Cephalosporin | Cefuroxime | 1 | 1.11 | 7 | 7.78 | 82 | 91.11 |
|  | Ceftriaxone | 19 | 21.11 | 4 | 4.44 | 67 | 74.44 |
|  | Ceftazidime | 13 | 14.44 | 6 | 6.67 | 71 | 78.89 |
|  | Cefotaxime | 6 | 6.67 | 9 | 10.00 | 75 | 83.33 |
|  | Cefepime | 20 | 22.22 | 2 | 2.22 | 68 | 75.56 |
| Fluoroquinolones | Ciprofloxacin | 30 | 33.33 | 6 | 6.67 | 54 | 60.00 |
|  | Levofloxacin | 15 | 16.67 | 10 | 11.11 | 65 | 72.22 |
| Tetracyclines | Doxycycline | 40 | 44.44 | 7 | 7.78 | 43 | 47.78 |
|  | Tetracycline | 19 | 21.11 | 6 | 6.67 | 65 | 72.22 |
| β-lactamase inhibitors | Piperacillin + Tazobactam | 25 | 27.78 | 7 | 7.78 | 58 | 64.44 |
|  | Amoxicillin-Clavulanate | 19 | 21.11 | 6 | 6.67 | 65 | 72.22 |
| Carbapenems | Imipenem | 24 | 26.67 | 9 | 10.00 | 57 | 63.33 |
|  | Meropenem | 25 | 27.78 | 1 | 1.11 | 64 | 71.11 |
| Folate inhibitors | Cotrimoxazole | 36 | 40.00 | 10 | 11.11 | 44 | 48.89 |
| Polymyxins | Colistin | - | - | 53 | 58.8 | 37 | 41.11 |

Comparative analysis between ICU and NICU isolates revealed consistently high resistance across both groups, with no statistically significant differences for most antibiotics. Notably, meropenem resistance was higher in ICU isolates (78.2%) than NICU isolates (63.6%) (p=0.07), while tetracycline resistance showed borderline significance (ICU 82.6% vs. NICU 61.3%, p=0.05) (Table 4). Hierarchical clustering and heatmap further demonstrated distinct resistance patterns, with β-lactams forming a dominant resistance cluster, while relatively susceptible agents such as colistin clustered separately (Figures 2C-D).

**Table 4:** Comparative antimicrobial susceptibility patterns of *Acinetobacter* spp. isolates from ICU (n = 46) and NICU (n = 44) patients.

| Antibiotics | ICU (n=46) |  |  | NICU (44) |  |  | p-value |
| --- | --- | --- | --- | --- | --- | --- | --- |
|  | S, n (%) | I, n (%) | R, n (%) | S, n (%) | I, n (%) | R, n (%) |  |
| Amikacin | 20 (43.4) | 7 (15.2) | 19 (41.3) | 23 (52.2) | 4 (9.0) | 17 (38.6) | 0.40 |
| Gentamycin | 9 (19.5) | 1 (2.1) | 36 (78.2) | 10 (22.7) | 5 (11.3) | 29 (65.9) | 0.71 |
| Cefuroxime | 1 (2.1) | 3 (6.5) | 42 (91.3) | 0 (0) | 4 (9.0) | 40 (90.9) | 0.32 |
| Ceftriaxone | 9 (19.5) | 1 (2.1) | 36 (78.2) | 10 (22.7) | 3 (6.8) | 31(70.4) | 0.71 |
| Ceftazidime | 8 (17.3) | 3 (6.5) | 35 (76.0) | 5 (11.36) | 3 (6.82) | 36 (81.8) | 0.41 |
| Cefotaxime | 2 (4.3) | 4 (8.70) | 40 (86.9) | 4 (9.09) | 5 (11.3) | 35 (79.5) | 0.36 |
| Cefepime | 9 (19.5) | 2 (4.3) | 35 (76.0) | 11 (25.0) | 0 (0) | 33 (75) | 0.53 |
| Ciprofloxacin | 13 (28.2) | 2 (4.3) | 31 (67.3) | 17 (38.6) | 0 (0) | 23(52.2) | 0.16 |
| Levofloxacin | 7 (15.2) | 4 (8.7) | 35 (76.0) | 8 (18.1) | 6 (13.6) | 30 (68.1) | 0.70 |
| Doxycycline | 16 (34.7) | 4 (8.7) | 26 (56.5) | 24 (54.5) | 3 (6.82) | 17 (38.6) | 0.05 |
| Tetracycline | 6 (13.0) | 2 (4.3) | 38 (82.6) | 13 (29.5) | 4 (9.09) | 27 (61.3) | 0.05 |
| Piperacillin + Tazobactam | 15 (32.6) | 3 (6.5) | 28 (60.8) | 10 (22.7) | 4 (9.09) | 30 (68.1) | 0.29 |
| Amoxicillin + Clavulanate | 10 (21.7) | 3 (6.5) | 33 (71.7) | 9 (20.4) | 3 (6.82) | 32 (72.7) | 0.88 |
| Imipenem | 12 (26.0) | 1 (2.17) | 33 (71.7) | 12 (27.2) | 8 (18.1) | 24 (54.5) | 0.89 |
| Meropenem | 9 (19.5) | 1 (2.17) | 36 (78.2) | 16 (36.3) | 0 (0) | 28 (63.6) | 0.07 |
| Cotrimoxazole | 19 (41.3) | 6 (13.0) | 21 (45.6) | 17 (38.6) | 4 (9.0) | 23 (52.2) | 0.79 |
| Colistin | - | 26 (56.5) | 20 (43.4) | - | 27 (61.3) | 17 (38.6) | 0.53 |

509 Here, intermediate susceptibility was categorized as resistant for statistical analysis. Statistical significance was defined as $p < 0.05$ .

**Table 5:** Distribution of carbapenemase and metallo-β-lactamase (MBL) genes in relation to ESBL phenotypes determined by phenotypic combined disc diffusion test (PCDDT).

| Genes | PCDDT + n (%) | PCDDT - n (%) | Odds Ratio (95% CI) | Fisher's exact test *p*-value |
| --- | --- | --- | --- | --- |
| *bla*<sub>NDM-1</sub> | 3 (10) | 9 (33.33) | 0.22 (0.06 - 0.87) | 0.0498 |
| *bla*<sub>IMP</sub> | 8 (26.67) | 0 (0) | ∞ (2.23 - ∞) | 0.0049 |
| *bla*<sub>VIM</sub> | 12 (40) | 20 (74.07) | 0.233 (0.07 - 0.68) | 0.0157 |
| *bla*<sub>OXA-23</sub> | 8 (26.67) | 19 (70.37) | 0.15 (0.04 - 0.46) | 0.0014 |
| *bla*<sub>OXA-58</sub> | 6 (20) | 2 (7.41) | 3.12 (0.66 - 16.11) | 0.258 |
| *blaAmpC* | 10 (33.33) | 3 (1.11) | 4 (1.06 - 14.67) | 0.061 |

Here, associations between gene prevalence and PCDDT status were evaluated using Fisher's exact test.
Statistical significance was defined as $p < 0.05$ ; \*\* $p < 0.01$ ; \* $p < 0.05$ ; ns, not significant.

### 3.3 Association of PCDDT Phenotype with Resistance Determinants

Among carbapenem-resistant isolates (n=55), 52.6% (n=30) were positive by phenotypic combined disc diffusion testing (PCDDT), showing positivity in the phenotypic combination disc diffusion test (PCDDT) (Figure 3A-B). PCDDT-positive isolates showed a higher prevalence of MBL-associated genes, particularly *blaVIM* (40%), followed by *blaIMP* (26.7%) and *blaNDM* (10%). Co-occurrence with class D carbapenemase genes (*blaOXA-23*, 26.7%; *blaOXA-58*, 20.0%) and AmpC-associated gene target (23.3%) were also observed. In contrast, PCDDT-negative isolates exhibited lower frequencies of *blaNDM* (33.3%), *blaOXA-58* (7.4%), and *blaAmpC* (1.1%), but higher prevalence of *blaVIM* (74.1%) and *blaOXA-23* (70.3%) (Table 4), suggesting alternative resistance mechanisms independent of phenotypically detectable MBL activity.

**Figure 3:**
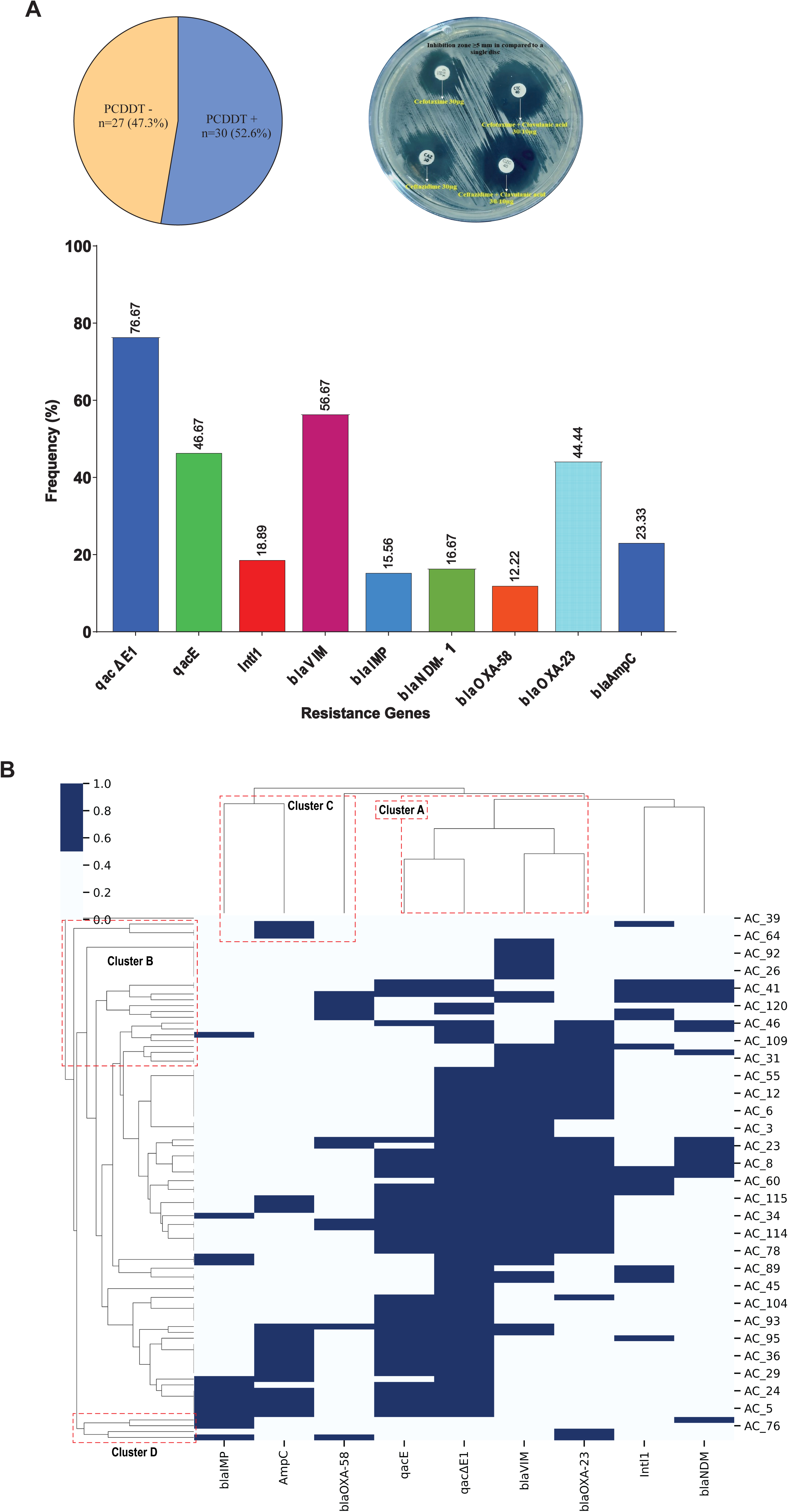
Phenotypic and genotypic profiling of resistance determinants in *Acinetobacter* spp. (A) Distribution of extended-spectrum β-lactamase (ESBL) production among carbapenem-resistant *Acinetobacter* isolates (n = 55), determined using the phenotypic combined disc diffusion test (PCDDT). ESBL positivity was defined by a ≥5 mm increase in the inhibition zone diameter for ceftazidime–clavulanic acid (CZC) or cefotaxime–clavulanic acid (CTC) compared with ceftazidime (CA), indicating clavulanate-mediated synergy with third-generation cephalosporins. Frequency distribution of resistance genes among *Acinetobacter* spp. isolates (N = 90). (B) Heatmap with hierarchical clustering of resistance genes across isolates, illustrating co-occurrence patterns based on the Jaccard similarity index and average linkage method.

### 3.4 Distribution of Resistance Genes

Biocide resistance determinants were highly prevalent, with *qac*Δ*E1* (76.6%) and *qacE* (46.6%) being the most frequently detected genes. Among β-lactam resistance determinants, *blaVIM* (56.6%) and *blaOXA-23* (44.4%) predominated. Other genes, including *blaAmpC* (23.3%), *IntI1* (18.8%), *blaNDM* (16.6%), *blaIMP* (15.5%), and *blaOXA-58* (12.2%), were detected at lower frequencies (Figure 3A). Hierarchical clustering revealed a dominant co-occurrence cluster comprising *qacE*, *qac*Δ*E1*, *blaVIM*, and *blaOXA-23*, representing the core multidrug-resistant genotype (Cluster A). Additional clusters represented genetically diverse subpopulations with variable resistance gene combinations, including *blaNDM*-driven profiles (Figure 3B).

### 3.5 Co-existence of Resistance Determinants

Gene co-occurrence analysis demonstrated significant associations between resistance determinants (Table 6). The most frequent pairing was *qac*Δ*E1* + *qacE* (n=32), showing strong correlation (p<0.001), high relative risk (RR=2.71), and the highest Jaccard index (0.604). Carbapenemase–biocide gene combinations were also prominent, including *qac*Δ*E1* + *blaVIM* (n=30; Jaccard=0.462) and *blaOXA-23* + *qac*Δ*E1* (n=27; Jaccard=0.458), both exhibiting significant associations. The strongest relative risk was observed for *blaOXA-23* + *blaVIM* (RR=3.05; p<0.001). Correlation analysis indicated moderate positive relationships between *qacE* and *qac*Δ*E1* (r=0.516) and between *blaOXA-23* and *blaVIM* (r=0.466), while negative associations were observed between *blaAmpC* and *blaVIM* (r=−0.366), among others (Figure 4A). Venn analysis of carbapenemase genes showed *blaVIM* as the dominant determinant (n=38), with limited co-occurrence among *blaIMP* and *blaNDM*. No isolate harbored all three genes simultaneously (Figure 4B).

**Figure 4:**
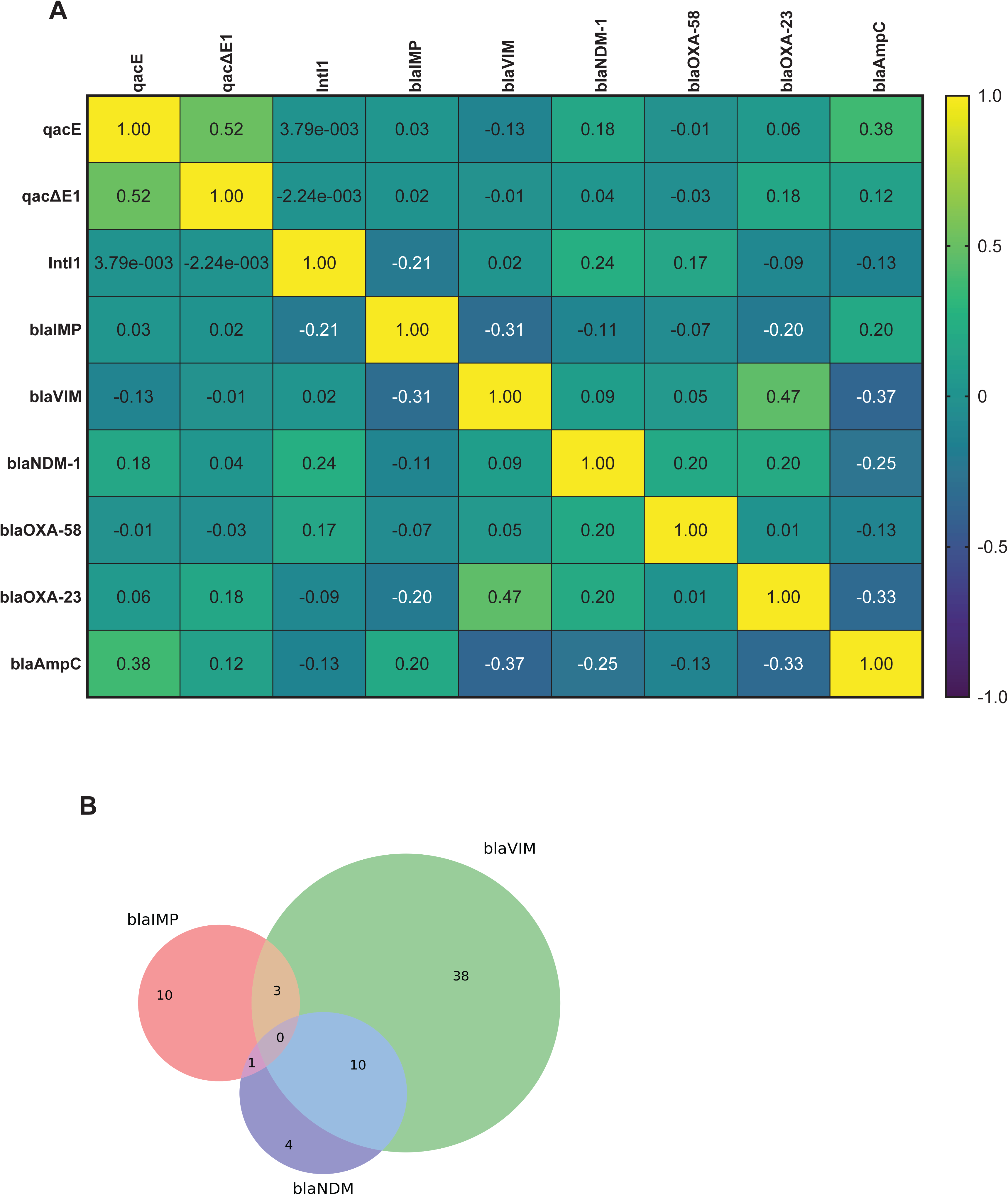
Association and co-occurrence analysis of resistance determinants in *Acinetobacter* spp. (A) Spearman correlation heatmap illustrating the co-occurrence relationships between β-lactamase genes and biocide (disinfectant) resistance determinants. (B) Co-existence patterns of carbapenemase genes among isolates, with Jaccard similarity analysis depicting the extent of shared gene profiles across *Acinetobacter* spp.

**Table 6:** Prevalence and co-occurrence of resistance gene pairs among *Acinetobacter* spp., including Jaccard similarity indices and relative risk estimates.

| Gene Pair | Co-existence (n) | Jaccard Index | *p*-value | Relative Risk (RR) |
| --- | --- | --- | --- | --- |
| *qacΔE1* + *qacE* | 32 | 0.604 | <0.001 | 2.71 |
| *qacΔE1* + *bla<sub>VIM</sub>* | 30 | 0.462 | 0.0256 | 1.46 |
| *bla<sub>OXA-23</sub>* + *bla<sub>VIM</sub>* | 27 | 0.563 | <0.001 | 3.05 |
| *bla<sub>OXA-23</sub>* + *qacΔE1* | 27 | 0.458 | <0.001 | 1.76 |
| *bla<sub>OXA-23</sub>* + *qacE* | 15 | 0.300 | 0.114 | 1.5 |
| *bla<sub>VIM</sub>* + *qacE* | 15 | 0.254 | 0.604 | 0.99 |
| *qacΔE1* + *bla<sub>NDM</sub>* | 12 | 0.214 | 0.066 | 1.44 |
| *Int11* + *qacΔE1* | 12 | 0.211 | 0.133 | 1.34 |
| *qacE* + *blaAmpC* | 10 | 0.313 | <0.001 | 3.59 |
| *qacE* + *bla<sub>NDM-1</sub>* | 10 | 0.27 | <0.001 | 2.24 |

### 3.6 Role of Class 1 Integron (IntI1) in Resistance Gene Dissemination

Network analysis identified IntI1 as a central node associated with the co-occurrence of multiple resistance determinants (Figure 5A). Strong associations were observed between *IntI1* and biocide resistance genes (*qacE*, *qac*Δ*E1*) as well as carbapenemase genes (*blaVIM*, *blaNDM*), with weaker links to *blaAmpC* and *blaOXA-58*. Among *IntI1*-positive isolates (n=17), *qac*Δ*E1* (76.5%) and *blaVIM* (58.8%) were the most prevalent co-harbored genes, followed by *qacE* (52.9%) and *blaNDM*/*blaOXA-23* (each 35.3%). *blaAmpC* was least frequent (11.8%) (Figure 5B). Intersection analysis revealed diverse gene combinations, including multideterminant profiles co-harboring biocide and carbapenemase genes. A subset of isolates carried complex resistance gene constellations, highlighting the role of integrons in facilitating multidrug resistance dissemination.

**Figure 5:**
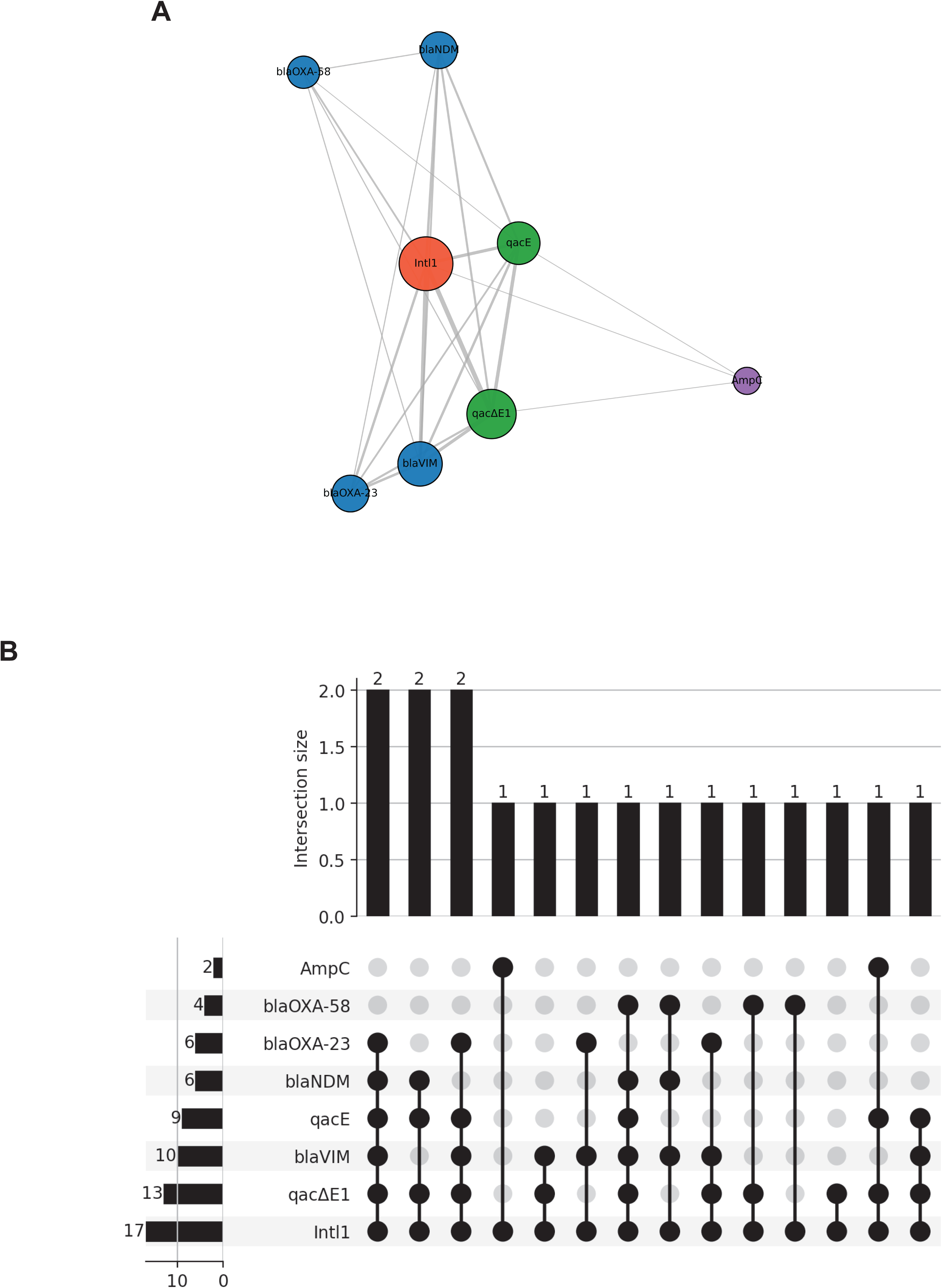
Network-based analysis of resistance gene interactions in *Acinetobacter* spp. (A) Co-occurrence network illustrating the central role of the class 1 integron (*intI1*) in linking carbapenemase genes (bla_NDM-1_, bla_VIM_, bla_OXA-23_, bla_OXA-58_), biocide resistance determinants (*qacE*, *qac*Δ*E1*), and *blaAmpC*. Node size corresponds to gene frequency, while edge thickness reflects the strength of co-occurrence between gene pairs. (B) Distribution and co-occurrence patterns of carbapenemase genes (bla_OXA-23_, bl_NDM-1_, bla_VIM_, bla_OXA-58_, *blaAmpC*) and biocide resistance determinants (*qacE*, *qac*Δ*E1*) in association with class 1 integrons (*intI1*) across *Acinetobacter*isolates.

## Discussion

The global emergence of *Acinetobacter baumannii* as a dominant nosocomial pathogen, particularly in intensive care settings, represents a critical threat to healthcare systems [3, 7, 22]. A substantial number of Acinetobacter bloodstream isolates were recovered from ICU (48.9%) and NICU (51.1%) patients during the study period, underscoring the organism’s pervasive role in severe hospital-acquired infections in Bangladesh. The exclusive recovery of isolates from bloodstream infections further emphasizes the clinical severity, with bacteremia serving as a key manifestation in critically ill and neonatal populations.

A striking finding was the high prevalence of multidrug resistance (MDR, 71.1%) and extensively drug resistance (XDR, 22.2%), consistent with the escalating resistance trends reported globally. These rates exceed or align with those reported across comparable ICU settings in South Asia and other regions, reflecting sustained antimicrobial selection pressure and inadequate infection control practices [7–9, 20, 23–26]. The comparatively higher XDR burden in ICU isolates suggests intensified antibiotic exposure and selective pressure in adult critical care environments, whereas prolonged hospitalization in NICUs appears to contribute to resistance accumulation.

The antimicrobial susceptibility profile revealed extensive resistance to β-lactams, including third- and fourth-generation cephalosporins (>74%) and carbapenems (>70%), indicating widespread therapeutic failure of frontline agents. This resistance is largely attributable to the production of β-lactamases, including carbapenem-hydrolyzing enzymes. Although colistin and amikacin retained partial activity, the observed decline in susceptibility is concerning, as these agents represent last-line therapeutic options. The emergence of reduced colistin susceptibility signals a narrowing therapeutic window and raises the risk of untreatable infections.

Molecular analysis further elucidated the genetic basis of resistance. The predominance of *blaVIM* (56.6%) and *blaOXA-23* (44.4%) highlights the dual contribution of metallo-β-lactamases and class D carbapenemases to carbapenem resistance. Notably, discrepancies between phenotypic detection (PCDDT) and genotypic findings underscore the limitations of conventional diagnostic assays, particularly for enzymes that are not inhibited by clavulanic acid. This phenotypic masking has important implications for surveillance, reinforcing the necessity of molecular diagnostics in high-risk clinical settings.

A key contribution of this study is the high prevalence of biocide resistance genes (*qac*Δ*E1* and *qacE*), which were strongly associated with carbapenemase genes. This co-occurrence suggests a potential link between disinfectant exposure and antimicrobial resistance selection. The presence of *qac* genes may indicate reduced susceptibility to quaternary ammonium compounds, although phenotypic biocide susceptibility was not evaluated in the present study. This finding has direct implications for infection control strategies, as it challenges the effectiveness of routine disinfection protocols [15, 17, 27, 28].

The role of mobile genetic elements was further evidenced by the detection of class 1 integrons (*IntI1*), which acted as central hubs connecting multiple resistance determinants. The observed associations between *IntI1* and resistance determinants are consistent with a potential role of class 1 integrons in resistance dissemination. However, direct evidence of gene linkage or horizontal transfer would require sequencing-based approaches. Such integron-mediated dissemination accelerates the evolution of highly resistant phenotypes and complicates containment efforts [13, 15, 18].

Co-occurrence and network analyses revealed complex resistance architectures, with frequent pairing of disinfectant and antibiotic resistance genes. The dominant *qac*Δ*E1–qacE* cluster and its association with carbapenemase genes highlight the emergence of multidimensional resistance profiles. These findings are compatible with the hypothesis of co-selection, although causality cannot be established from the present data [15, 26].

From a clinical and public health perspective, these findings have several critical implications. First, the high burden of MDR/XDR *Acinetobacter* infections severely limits therapeutic options, increasing reliance on toxic or less effective agents. Second, the coexistence of disinfectant and antibiotic resistance genes threatens the effectiveness of infection prevention measures. Third, the integron-mediated dissemination of resistance underscores the need for genomic surveillance to track transmission dynamics [11, 26].

Several methodological limitations should be considered. Species-level identification was not performed, and therefore the distribution of resistance determinants among individual *Acinetobacter* spp. could not be assessed. Furthermore, molecular typing approaches such as MLST, PFGE, rep-PCR, or whole-genome sequencing were not available, preventing assessment of clonal relatedness, outbreak potential, and genetic linkage between resistance determinants and class 1 integrons.

## Conclusion

Bloodstream infections caused by multidrug- and extensively drug-resistant *Acinetobacter* spp. are highly prevalent in ICU and NICU settings, severely constraining treatment options. The frequent co-occurrence of carbapenemase genes, qac determinants, and class 1 integrons highlights the complex resistance landscape among Acinetobacter bloodstream isolates. These findings highlight an urgent need for integrated antimicrobial stewardship, molecular surveillance, and reassessment of disinfection practices to effectively control transmission in critical care environments.

## Acknowledgement

We sincerely thank the legal guardians of patients in the ICU and NICU for providing informed consent for participation in this study. We also acknowledge the participating hospitals for their cooperation. In addition, we express our gratitude to the laboratory members of the Department of Pathology and Parasitology, Chattogram Veterinary and Animal Sciences University (CVASU), for their technical assistance and support throughout the study.

## Ethical approval

This study was conducted in accordance with the ethical standards of the relevant institutional and national research committees. Ethical approval was obtained from the Institutional Review Committee (IRC) of Chattogram Veterinary and Animal Sciences University (CVASU). Written informed consent was obtained from the legal guardians of all participants prior to sample collection.

## Author Contributions

T.M.R., F.F.B.H., P.A.J.: Conceptualization, Investigation, Methodology, Software, Formal Analysis, Writing – Original Draft Preparation; S.C., S.M., S.A., A.A.K., S.M., M.A., N.A.: Data Curation, Methodology, Validation, Visualization, Writing – Review & Editing; T.M.R., A.M.A.M.Z.S.: Conceptualization, Supervision, Writing – Review & Editing, Resources, Project Administration, Funding Acquisition, Writing – Review & Editing

## Data Availability

All data generated and analyzed during this study are included in this published article and presented in the main text, figures, and tables. Details of all primers used for molecular analyses, including their sequences and sources, are provided in Table 1. Additional supporting information may be made available by the corresponding author upon reasonable request, subject to institutional regulations, ethical approval conditions, and participant consent regarding data sharing.

